# *Cis* mutagenesis *in vivo* reveals extensive noncanonical functions of Dscam1 isoforms in neuronal wiring

**DOI:** 10.1101/2022.04.14.488281

**Authors:** Shixin Zhang, Xi Yang, Haiyang Dong, Bingbing Xu, Lili Wu, Jian Zhang, Guo Li, Pengjuan Guo, Lei Li, Ying Fu, Yiwen Du, Yanda Zhu, Feng Shi, Jianhua Huang, Haihuai He, Yongfeng Jin

## Abstract

*Drosophila Dscam1* encodes ten thousand of cell recognition molecules via alternative splicing, which is required for nervous function. However, the underlying mechanism has not fully understood. Here we revealed extensive noncanonical functions of Dscam1 isoform diversity in neuronal wiring. We generated a series of allelic *cis* mutations in Dscam1, encoding normal number of isoforms albeit with altered isoform composition. Despite normal dendritic self-avoidance and self/non-self discrimination in da neurons, which is consistent with canonical self-avoidance model, these mutants exhibited strikingly distinct spectra of phenotypic defects in three classes of neurons: up to ∼60% defects in mushroom bodies, a significantly increased branching and growth in da neurons, and mild axonal branching defects in mechanosensory neurons. These data suggest that proper Dscam1 isoform composition is required for axon and dendrite growth in diverse neurons. This unprecedented splicing-tuned regulation unlocks new insights into the molecular mechanisms that Dscam1 isoform diversity is engaged in.

## Introduction

Developing neurons interact in specific and stereotyped ways to form the neural circuits that underlie complex brain function.(Sanes and Zipursky, 2020) Over the past 20 years, numerous gene families have been implicated in target recognition, including fly *Dscam1* and vertebrate clustered *Pcdhs*.(Schmucker et al., 2000; Wu and Maniatis, 1999) *Drosophila Dscam1* encodes 38,016 isoforms through alternative splicing of four exon cluster of exon 4, 6, 9, and 17, each comprising one of 19,008 alternative ectodomains linked to one of two alternative transmembrane domains. Identical Dscam1 ectodomains interact strongly and mediate homophilic repulsion, but interactions between different ectodomains are weak.(Wojtowicz et al., 2007) Each neuron produces only a small subset of Dscam1 isoforms (10-50 isoforms) in a largely stochastic manner, endows each individual neuron with a unique molecular identity for self-recognition in the nervous system.(Miura et al., 2013; Neves et al., 2004) This pattern results in repulsion between neurites of a single neuron, while allowing contact with other neurons of the same type. Genetic studies showed that thousands of isoforms are required to distinguish between self and non-self during self-avoidance, and for normal patterning of axons and dendrites.(Hattori et al., 2009; Hattori et al., 2007) In this case, there is no requirement for specific isoforms; it is only important that the subset of isoforms expressed by one neuron are different from those of others.(Chen et al., 2006; Hattori et al., 2009; Hattori et al., 2007; He et al., 2014; Liu et al., 2020; Sanes and Zipursky, 2020; Wang et al., 2004; Wu et al., 2012; Zhan et al., 2004; Zipursky and Sanes, 2010). While these mechanisms have been shown to underlie spacing of mechanosensory neuron dendrites and mushroom body development, it remains unclear how general this self-avoidance model explain Dscam1 functions in neural circuit assembly.

However, these functional experiments using RNAi knockdown, knock-ins, or genomic deletions have either resulted in changed overall protein levels, a reduced isoform diversity, or both. Therefore, it remains unclear whether these defective phenotypes observed in mutants are caused by changes in the Dscam1 protein levels, the isoform number, or both. Importantly, reducing Dscam1 isoform diversity led to phenotypic defects not only resulting from loss of function, but also gain of function. A definitive strategy to address the noncanonical functions of Dscam1 diversity would construct mutant flies with identical protein levels as wild type controls, and with thousands of isoforms, which would be sufficient to confer the normal self-avoidance and self/non-self discrimination. If these mutant flies exhibit an obvious phenotypic defect, these defects should be attributed to noncanonical role of Dscam1 isoforms rather than self-avoidance.

In this study, we used *in vivo* mutagenesis to reveal that variable exon 9 selection is mediated by bidirectional docking site-selector base pairing. Interestingly, this led us to construct mutant flies encoding a comparable number of Dscam1 isoforms as WT albeit with altered isoform composition. These mutants exhibited normal dendritic self-avoidance and self/non-self discrimination in da neurons, which is consistent with canonical self-avoidance model. Surprisingly, these mutants exhibited strikingly distinct spectra of phenotype defects in three classes of neurons: up to ∼60% defects in mushroom bodies, a significantly increased branching and growth in da neurons, and mild axonal branching defects in mechanosensory neuron. These data suggest that Dscam1 isoform composition is required for normal growth in diverse neurons.

## Results

### Construction of mutant flies with normal Dscam1 overall protein levels and an identical isoform number but with altered Ig7 composition

The initial motivation for our studies was to explore the molecular mechanism underlying alternative splicing of exon cluster 9 in *Drosophila Dscam1*. We have previously identified the docking site in the intron upstream of exon 10 of *Drosophila Dscam1*, which may pair with the selector sequence downstream of only a few exon 9 variants.(Hong et al., 2021; Yang et al., 2011) Considering the presence of dual docking sites and bidirectional base pairings in the exon 9 cluster of distantly related lepidopteran, coleopteran and hymenopteran *Dscam1* (Figure S1A), (Dong et al., 2021; Yue et al., 2016) we speculated that an analogous docking site may be located in the intron upstream exon 9.1 of *Drosophila Dscam1*. However, sequence alignment revealed only relatively limited conservation across *Drosophila* (Figure S1B). Since these introns are only ∼150 nt in size, we speculated that an analogous upstream docking site, if it exists, should be in the middle region of intron 8. To address this issue, we used CRISPR/Cas9 technology to delete the middle intronic region and then investigate the effect on the inclusion of exon 9 variants. We generated a series of homozygous viable flies varying in the degree of deletion (designated *Dscam1*^Δ9uD1-Δ9uD5^) (Figure 1A). We have not observed obvious phenotypic defects in all *Dscam1*^Δ9uD1-Δ9uD5^ adults. Reverse transcription PCR (RT-PCR) analysis indicated that there was no significant difference in the overall inclusion of exon 9 in the head and different developmental stages of *Dscam1*^Δ9uD^ compared with wild-type (Figure 1B). Western blotting analysis showed that the Dscam1 level was indistinguishable between *Dscam1*^Δ9uD^ and wild-type (Figure 1C). These data indicated that *Dscam1*^Δ9uD^ mutant flies displayed normal Dscam1 expression levels as wild-type controls.

**Figure 1.**
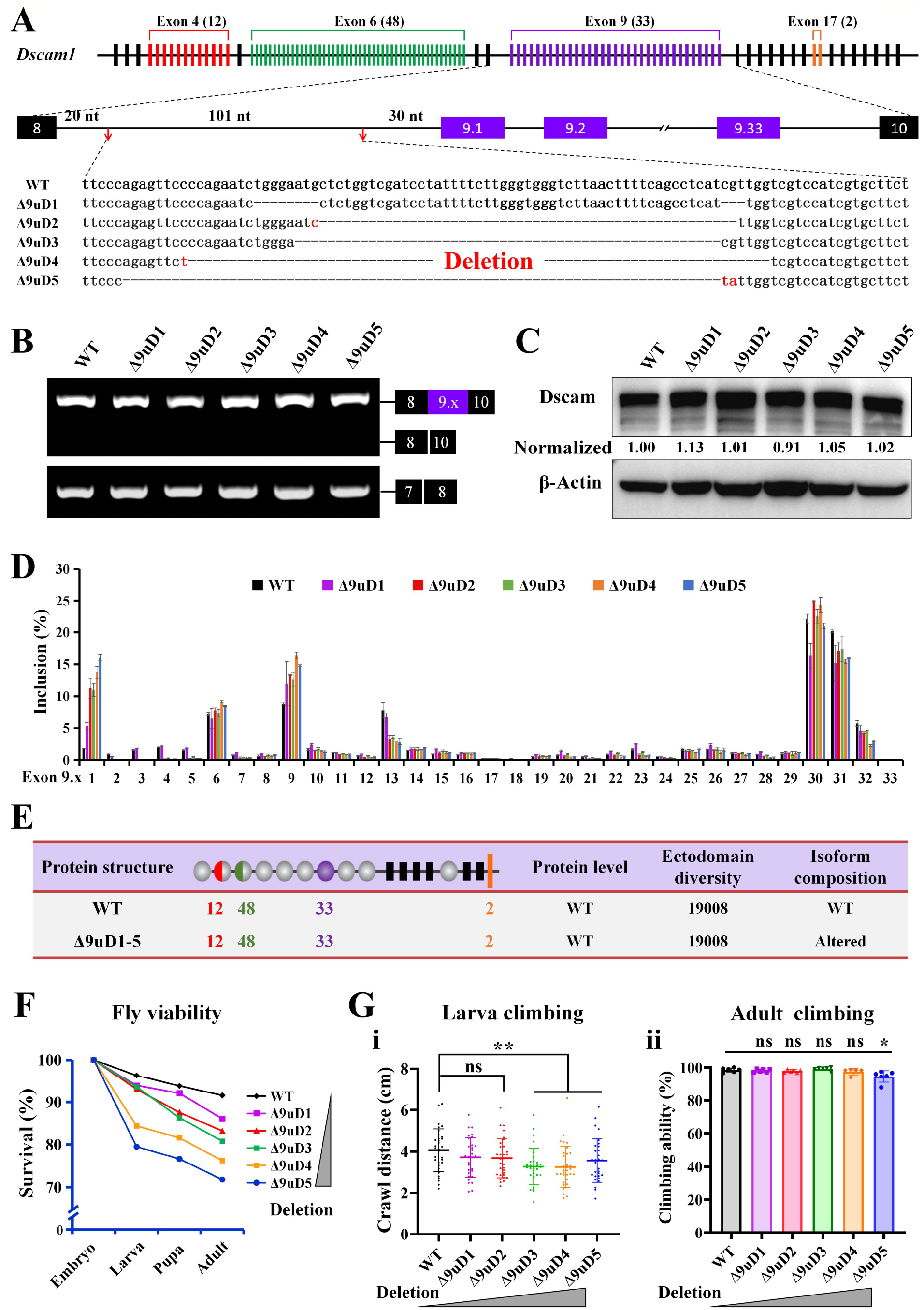
Construction and characterization of *Dscam1* isoform composition-altered mutants. (see also Figures S1, 2) **(A)** Schematic of the *cis* regulatory region targeted by CRISPR-Cas9. The red arrow indicates the position of the designed sgRNAs. The dashed lines represent the deleted sequences. **(B)** RT-PCR analysis showed that the overall inclusion of exon 9 is not affected. **(C)** Western blotting analysis showed that the Dscam1 level of *Dscam1*^Δ9uD^ was indistinguishable from those of wild type control. **(D)** *Dscam1*^Δ9uD^ mutants show mild to global alteration of expression of individual exon 9 variant compared with wild type control. **(E)** Characterization of *Dscam1*^Δ9uD^ compared with WT. **(F)** Survival rates of wild-type and *Dscam1*^Δ9uD^ mutants during development. **(G)** Effect of changed expression of exon 9 variants on fly locomotion. Climbing ability of mutant larva (i) and adult (ii) was tested. ns, not significant; *, P < 0.05; **, P < 0.01; (Student’s t-test, two-tailed).

To determine how the frequency of individual exon 9 variants was changed in *Dscam1*^Δ9uD^ mutants, we examined the relative usage of exon 9 variants in the RT-PCR products containing exon 9 using high-throughput sequencing. Analysis of sequencing revealed a profound change in the frequency of exon 9 variants in the *Dscam1*^Δ9uD^ flies (Figure 1D). *Dscam1*^Δ9uD1^ mutant with the smallest deletion exhibited subtle change of frequency of exon 9s, while *Dscam1*^Δ9uD2-Δ9uD5^ with the larger deletion exhibited global change in exon 9 inclusion. In addition, we observed a shared and specific change of expression at different developmental stages of mutant flies (Figure S2A–D). Notably, except for a few exon 9s, this pattern of change in the frequency of most exon 9 variants in *Dscam1*^Δ9uD4^ mutants was largely in contrast with the changing pattern in *Dscam1*^Δ9D^ mutants, which lack the downstream docking site (Figure S2E).(Hong et al., 2021) Thus, this splicing outcome of these mutant flies seems consistent with the regulatory model by dual docking sites and bidirectional base pairings. Interestingly, this led us to construct mutant flies encoding a comparable number of Dscam1 isoforms as WT albeit with global changes in exon 9 variant composition (Figure 1E). These viable homozygous mutants allowed us to explore the role of Dscam1 isoform composition in fly development and neuronal wiring. Below we focus to assess phenotype defects in three classes of neurons of *Dscam1*^Δ9uD1-Δ9uD5^ mutants.

### Altered composition of Ig 7 variants causes subtle to mild fly viability and locomotion defects

We first investigated the effect of deletion on the survival from embryo to adult stage in *Dscam1*^Δ9uD1-Δ9uD5^ mutant flies. The survival rates of *Dscam1*^Δ9uD1-Δ9uD5^ embryo were mildly decreased with the increase of deletion (Figure 1F). *Dscam1*^Δ9uD1^ mutants exhibited subtle survival defect, but *Dscam1*^Δ9uD4^ and *Dscam1*^Δ9uD5^ mutants exhibited ∼75% survival rate compared with ∼90% of the wild type control (Figure 1F). The difference in survival rate is mainly reflected in the embryo hatching stage, but the pupation rates and eclosion rates were no significant difference. This data suggests that altered Ig 7 isoform composition is detrimental to embryonic development.

To analyze the effect of Ig 7 isoform composition alteration on fly behavior, we carried out a crawling ability of the *Dscam1*^Δ9uD1-Δ9uD5^ mutant larvae. We observed a subtle to a mild decrease in the crawling ability of fly mutant larvae with the increase of deletion (Figure 1G, panel i). In addition, there was no significant difference in locomotion rate between *Dscam1*^Δ9uD1-Δ9uD4^ and wild-type adults, except for *Dscam1*^Δ9uD5^ with the largest deletion (Figure 1G, panel ii). Taken together, these data indicated that the proper composition of Dscam1 Ig7 variants was required for normal fly viability and locomotion.

### *Dscam1*^Δ9uD^ exhibit normal dendritic self/non-self discrimination but an obvious increased branching and growth in da neurons

To examine a potential role of Dscam1 diversity in dendrite wiring, we first compare dendritic morphology between *Dscam1*^Δ9uD^ mutants and WT in da neuron. *Dscam1*^Δ9uD^ mutants have no obvious differences in their characteristic overall shape or the dendritic territory borders, as compared with controls (Figure 2A). As expected, self-dendrites of class I dendritic arborization neurons in *Dscam1*^Δ9uD1-Δ9uD5^ mutants avoided each other as in wild-type controls, whereas *Dscam1*^null^ mutants exhibited severe overlap and fasciculation of self-dendrites (Figure 2A, B). Quantitative analysis showed that class I da (vpda) neurons in *Dscam1*^Δ9uD2-Δ9uD5^ mutants exhibited a slightly more overlap between dendritic self-branches than WT (Figure 2B). In addition, similar dendritic self-repulsion has been observed in the other two types of class I neurons (ddaD and ddaE) of *Dscam1*^Δ9uD2-Δ9uD5^ mutants (Figure 3B, C, F). Furthermore, we found that the number of overlaps between class I (vpda) and class III (v′pda) dendrites are not significantly different among various *Dscam1*^Δ9uD1-Δ9uD5^ mutants and wild-type control (Figure 2C; Figure S4). These results suggest that the number of Dscam1 isoforms in *Dscam1*^Δ9uD1-Δ9uD5^ mutants was sufficient to distinguish between self and non-self neurites.

**Figure 2.**
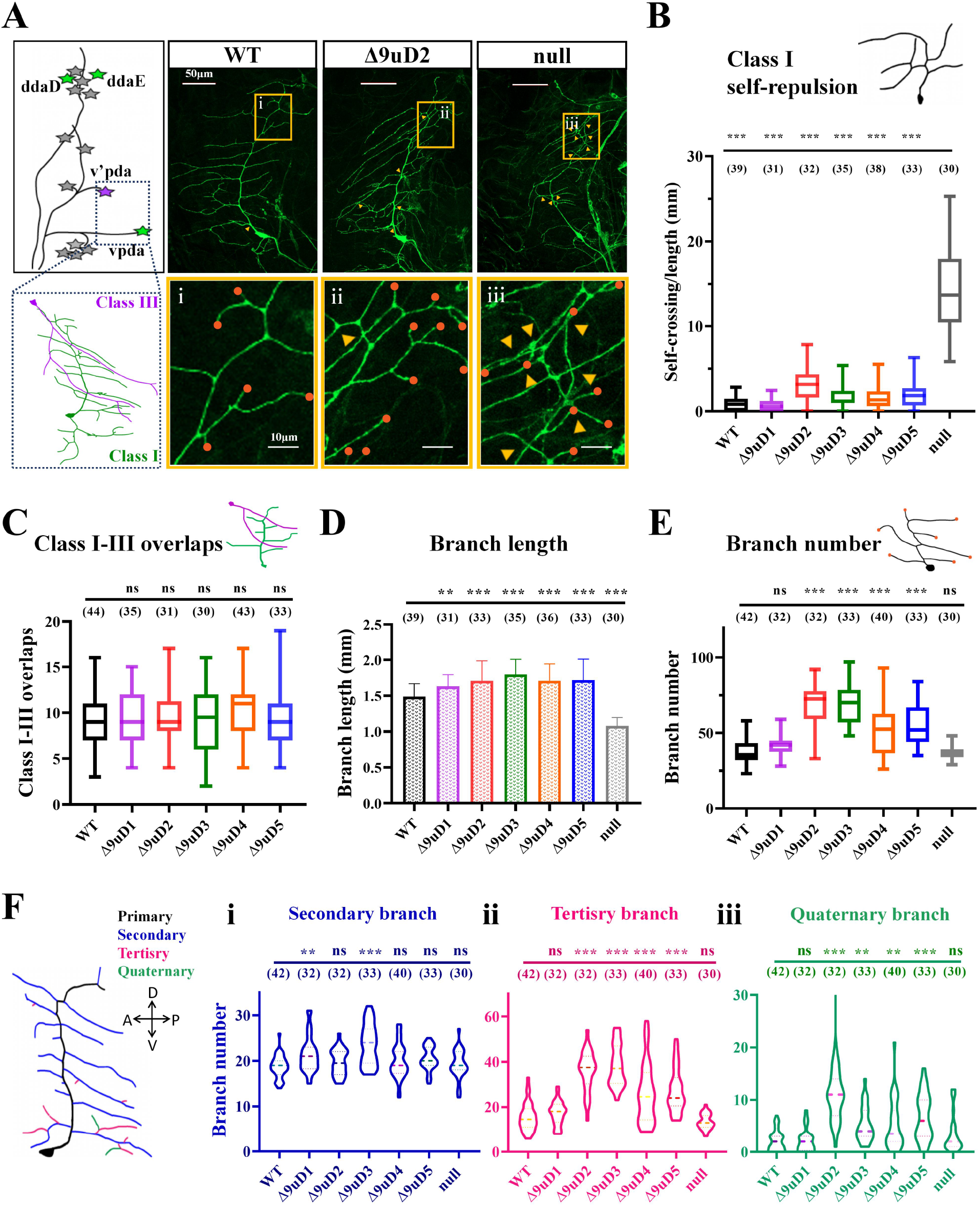
*Dscam1*^Δ9uD2-Δ9uD5^ mutants showed an increased branching in class I da neuron. (see also Figures S3, 4) **(A)** Schematic of an abdominal hemisegment of dendritic arborization (da) neurons. Class I da neuron are shown to green while class III are shown to magenta. Representative images of dendritic arborization neuron dendrites in different *Dscam1* mutants are shown. Scale bar, 50μm. Yellow boxes indicate detailed views in (i,ii,iii) to illustrate *Dscam1*^Δ9uD2-Δ9uD5^ show a slight increase in self-crossing compared to *Dscam1*^null^ and a marked increase in the number of branches. Scale bar, 20μm. Yellow arrows indicate points of self-crossing between sister branches, orange points represent branches in selected area. **(B)** *Dscam1*^Δ9uD2-Δ9uD5^ vpda show slight self-crossing while *Dscam1*^Δ9uD1^ were normal compared with wild-type. **(C)** The number of overlaps between class I and class III dendrites were indistinguishable between *Dscam1*^Δ9uD1-Δ9uD5^ mutants and wild-type controls. **(D)** Total dendritic length of *Dscam1*^Δ9uD2-Δ9uD5^ was more than those of wild-type neurons. **(E)** Vpda shows a significant increase in total branches in *Dscam1*^Δ9uD2-Δ9uD5^. **(F)** Vpda shows an increase in branches in *Dscam1*^Δ9uD2-Δ9uD5^. Vpda falls into four branches, the primary branch is the limb (color in black), the secondary branch grows directly from the middle primary branch (color in blue), the tertiary branch is the small side branch on the secondary branch (color in pink) and the quaternary branch is the other smaller branch (color in green). The tertiary branches and quaternary branches are increased observably in *Dscam1*^Δ9uD2-Δ9uD5^. Numbers in parenthesis are neurons analyzed. Data in boxplot are represented as median (dark line), 25%-75% quantiles (box), and error bar (1.5× quartile range). *, P < 0.05; **, P < 0.01; ***, P < 0.001; ns, not significant (Student’s t-test, two-tailed).

**Figure 3.**
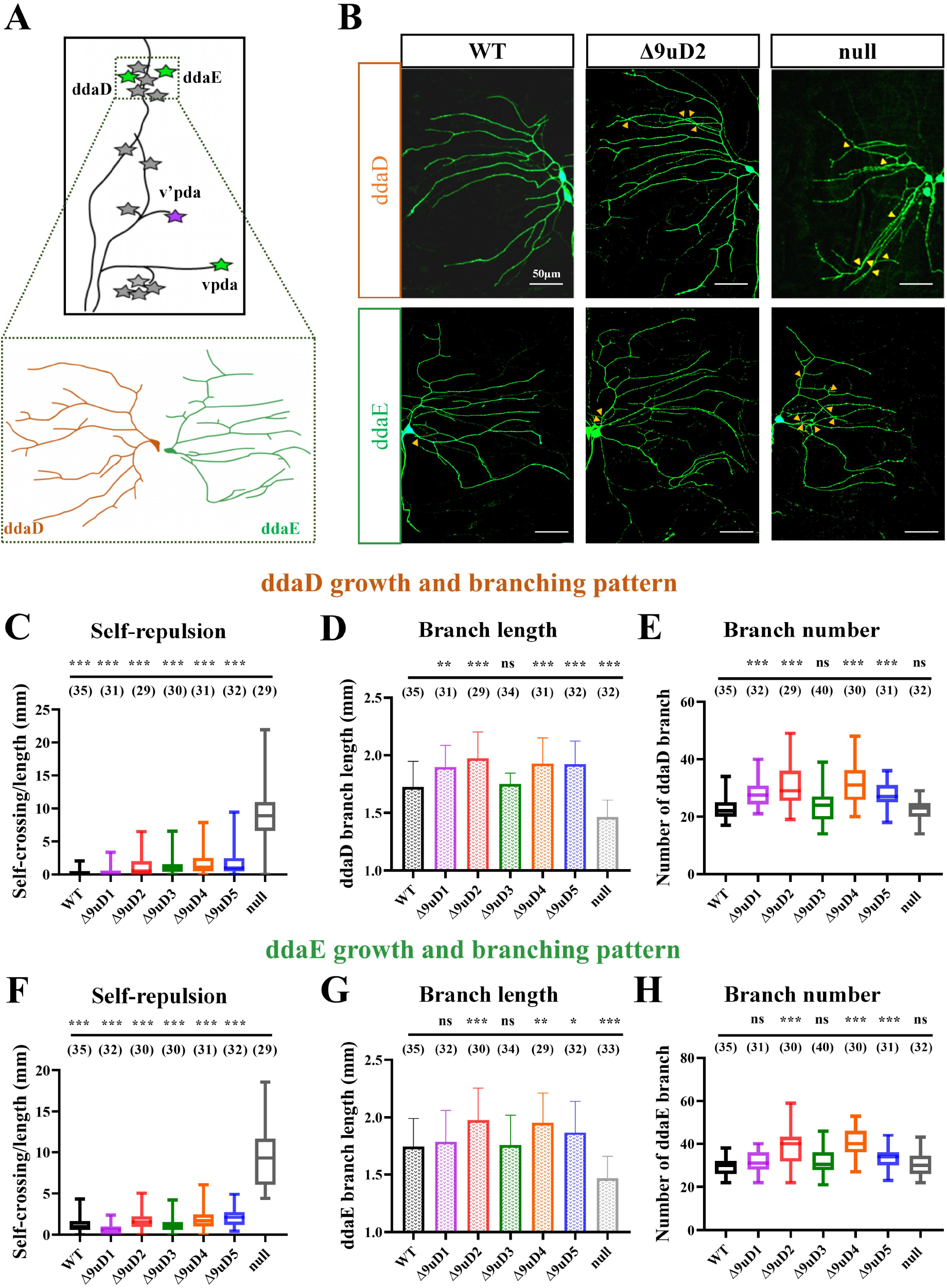
Class I da neurons exhibit dendritic patterning defects. (see also Figure S3) **(A)** Relative positions of ddaD and ddaE and their branching patterns. **(B)** Representative images of ddaD and ddaE dendrites in different *Dscam1* mutants. All neurons were visualized with GFP and their corresponding color-coded tracings are shown below. **(C-E)** The branching pattern of ddaD in the *Dscam1*^Δ9uD^ mutant third-instar epidermis changes. **(c)** The self-crossing of ddaD in the *Dscam1*^Δ9uD^ mutants are slightly increased. **(d)** Dendritic length of *Dscam1*^Δ9uD2-Δ9uD5^ are increased compared to wild-type neurons and *Dscam1*^null^ mutants. **(e)** ddaD shows a significant increase in total branches in *Dscam1*^Δ9uD^. **(F-H)** ddaE also show same branching pattern in the *Dscam1*^Δ9uD^ mutant third-instar epidermis. Numbers in parenthesis denote da neurons analyzed for each mutant. ns, not significant; *, P < 0.05; **, P < 0.01; ***, P < 0.001 (Student’s t-tests, two-tailed).

We next compared the dendritic length and branch number between *Dscam1*^Δ9uD^ mutants and wild-type control. Quantification revealed that the total dendritic length of *Dscam1*^Δ9uD2-Δ9uD5^ mutants was significantly more than that of wild-type neurons while *Dscam1*^Δ9uD1^ mutants exhibited a slight increase (Figure 2D). *Dscam1*^Δ9uD2-Δ9uD5^ mutants exhibited significant increases of total dendritic branch numbers by ∼30–100% (Figure 2E). In particular, the total dendritic branch per neuron has risen up to 70 in *Dscam1*^Δ9uD2^ mutants, which is nearly two-fold of 37 in wild-type controls. Further analysis revealed that these increases were caused by additional tertiary and quaternary branches, but primary and secondary dendrites were not largely affected (Figure 2F), indicating that Dscam1 likely involved new dendritic branch formation. Since *Dscam1*^Δ9uD^ alleles express the same overall expression level and encode a comparable number of Dscam1 isoforms as wild-type controls, we therefore conclude that these increased branching should be attributed to the altered composition of Dscam1 Ig7 variants.

Similarly, we have observed a notable increase in the total dendritic length of ddaD and ddaE neurons in *Dscam1*^Δ9uD2-Δ9uD5^ mutants compared with WT (Figure 3D, G). *Dscam1*^Δ9uD2-Δ9uD5^ mutants exhibited significant increases in the total number of dendritic branches in ddaD and ddaE neurons (Figure 3E, H). Notably, there are considerable differences in length and number of dendrite branches among different *Dscam1*^Δ9uD^ mutants (Figures 2, 3), suggesting that specific Dscam1 repertoire of each cell may intricately function in dendrite formation. Taken together, our results indicated that Dscam1 isoforms are not only required for discrimination between self and non-self neurites, but also for normal dendrite growth and branching in *Drosophila* da neuron. The former is no requirement for specific isoforms, while the latter involved specific isoforms, at least for Ig7 variants.

### *Dscam1*^Δ9uD4^ and *Dscam1*^Δ9uD5^ exhibit >50 % mushroom body defects

We next assessed whether and how the altered composition of Ig 7 variants caused mushroom body (MB) morphological defects. Unexpectedly, we observed 15-67% MB phenotype defects in *Dscam1*^Δ9uD2-Δ9uD5^ mutant flies. MB phenotype improved as the deletion size reduced (Figure 4B). In particular, we found that more than 67% and 56% of brains displayed MB phenotype defects in *Dscam1*^Δ9uD4^ and *Dscam1*^Δ9uD5^ mutant flies, respectively (Figure 4B). Moreover, transheterozygous mutants *Dscam1*^Δ9uD4^*/Dscam1*^Δ9uD5^ show abnormal MB phenotypes in 50% of brains, which is close to *Dscam1*^Δ9uD4^ and *Dscam1*^Δ9uD5^ mutants (Figure 4B). All *Dscam1*^Δ9uD2-Δ9uD4^*/Dscam1*^*+*^ heterozygous mutants exhibited < 5% of MB phenotype defect (Figure 4B). Mushroom body defect phenotypes included absence, truncating, thinning, and mis-projection of the lobes (Figure 4A, B).

**Figure 4.**
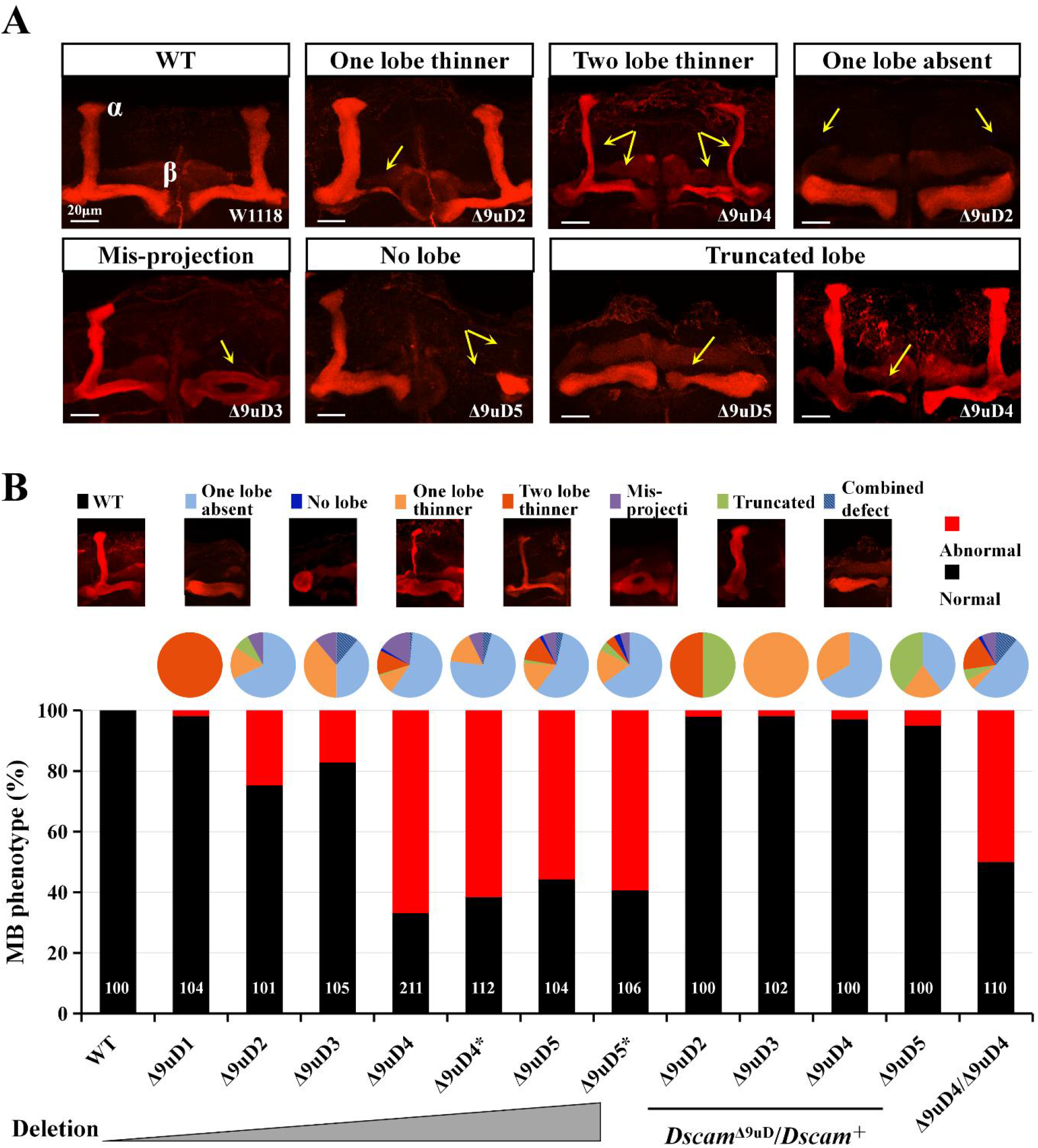
*Dscam1*^Δ9uD^ mutants exhibit mushroom body defects. (see also Figure S5) **(A)** Representative images of mushroom body (MB) lobe defects in mutant animals. Anti-FasII staining visualizing the α and β lobes of mushroom bodies in the adult brain; yellow arrows indicate lobe defects. **(B)** MB phenotypes of *Dscam1*^Δ9uD^ mutant flies. The pie charts above the bar graph display the proportion of different types of MB defects in different mutants, respectively. Numbers in parentheses represent the numbers of brain hemisphere examined for each genotype. *Dscam1*^Δ9uD4*^ and *Dscam1*^Δ9uD5*^, which lacked the same sequence as *Dscam1*^Δ9uD4^ and *Dscam1*^Δ9uD5^, were independently generated via homologous recombination.

To preclude the possibility of defects caused by the off-target effect, we used homologous recombination to independently construct two homozygous mutants (designated *Dscam1*^Δ9uD4*^ and *Dscam1*^Δ9uD5*^), which lacked the same sequence as *Dscam1*^Δ9uD4^ and *Dscam1*^Δ9uD5^, respectively (Figure S5). These *Dscam1*^Δ9uD4*^ and *Dscam1*^Δ9uD5*^ mutants exhibited a comparable spectrum of MB phenotype defects as *Dscam1*^Δ9uD4^ and *Dscam1*^Δ9uD5^ mutants, respectively (Figure 4B). We therefore conclude that such high penetrance of the MB defective phenotypes should be attributed to the globally altered composition of Dscam1 Ig7 variants. Since *Dscam1*^Δ9uD^ alleles express the same overall expression level and encode a comparable number of Dscam1 isoforms as wild-type controls, such high penetrance of the MB defective phenotypes cannot be explained by the canonical self-avoidance model.

### Altered composition of Ig 7 variants causes extensive MB axonal growth and branching defects in single-cell clones

To elucidate how the MB defective phenotypes of in *Dscam1*^Δ9uD^ mutant were correlated with MB axons, we used MARCM analysis to assess the axonal phenotypes at single-cell resolution.(Lee and Luo, 1999) In contrast wild-type control clones (n = 29), more than 40% of *Dscam1*^Δ9uD^ mutant neurons (n > 30) exhibited either a growth defect (the formation of shortened branches than in wild-type), or a branching defect, or a guidance defect, or sometimes a combination of them (Figure 5A,B). The whole MB defects were strongly correlated with single clone defect (Figure 5C). Axon growth was most sensitive to the change of Ig 7 variants of Dscam1. Axonal growth defects which mostly exhibited the shortened axon branches, which accounted for up to ∼55% of all defects in *Dscam1*^Δ9uD2-Δ9uD5^ (Figure 5D). This axonal shortening is consistent with the presence of truncated lobes (Figure 4B), suggesting the presence of truncated lobes might be caused by axon truncation.

**Figure 5.**
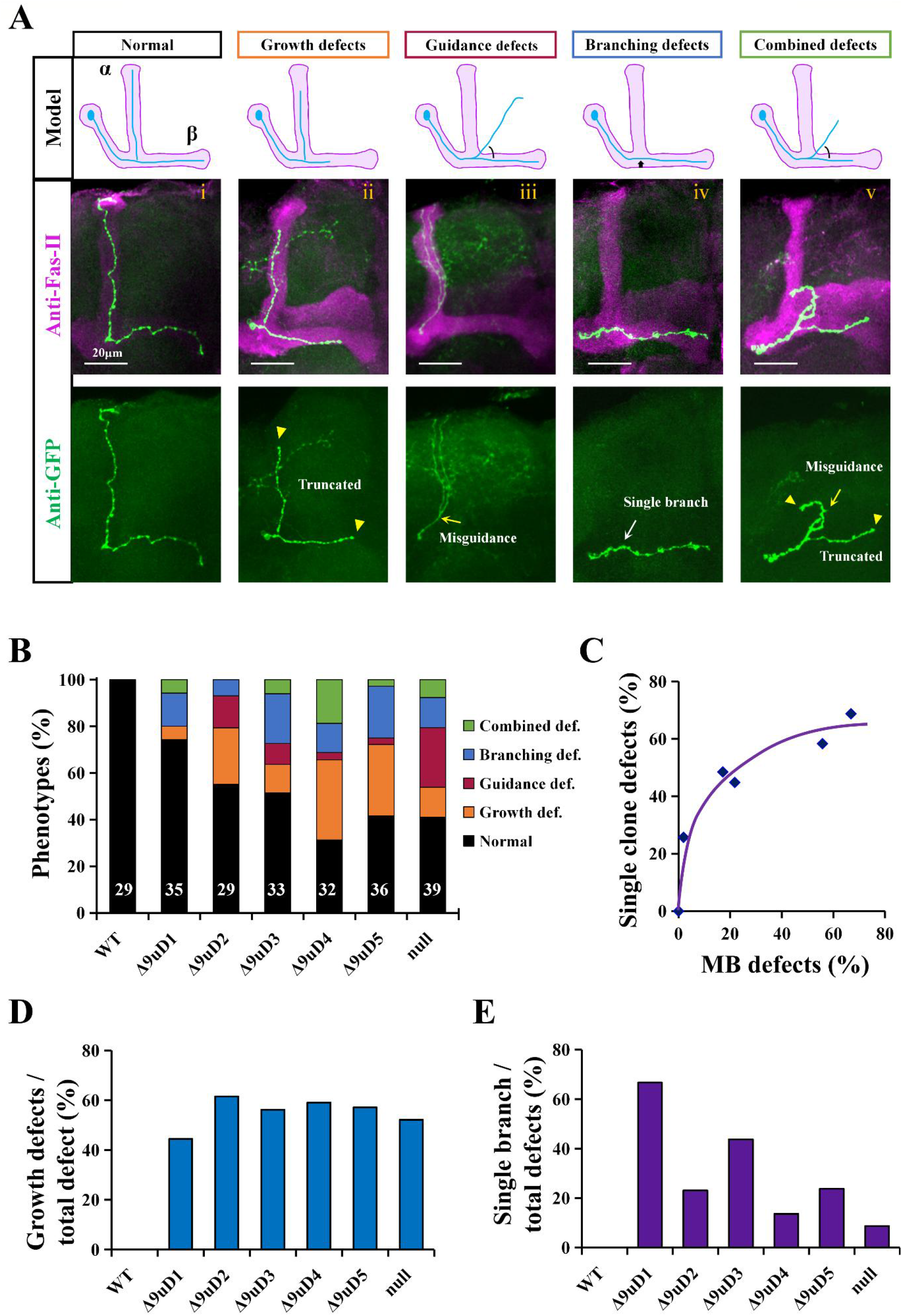
Phenotypic analysis of *Dscam1*^Δ9uD^ mutant mushroom body axons at single-cell clones. **(A)** Common types of mushroom body monoclonal development defects. All drawings were of adult mushroom body of the brain hemisphere. Top, model of different defects. Middle and bottom, representative images of single-cell clone defects. Anti-FasII staining visualizing the whole mushroom bodies (magenta) while single-cell clones showed green (Anti-GFP). Middle, i: wild type. ii: growth defect. iii: guidance defect. iv: single branch. v: multiple branch. vi: combined defect which show both growth and guidance defect. Bottom, yellow arrowhead indicates truncated branch, yellow arrow shows misguidance branch, white arrow identifies the abnormal branch points. **(B)** Quantification of mushroom body axon defects in single-cell clones. Numbers in parenthesis denote single-clones analyzed for each mutant. **(C)** The whole MB defects were strongly correlated with single clone defect. **(D)** *Dscam1*^Δ9uD1-Δ9uD5^ exhibited the high proportion of axonal growth defects. **(E)** *Dscam1*^Δ9uD1-Δ9uD5^ exhibited the high proportion of single axonal branches.

The axon branching also sensitivity to the change of Ig 7 variants, accounting for up to ∼40% of the axonal defects (Figure 5B). Branching defects included the generation of α-but not β-axonal branches without axon bifurcation at the peduncle end and vice versa (Figure 5A). About 10–40% of single neurons exhibit such defects in *Dscam1*^Δ9uD1-Δ9uD5^ (Figure 5E). Single branch can explain the absence and thinning phenotype in MB, even no lobe when simultaneous growth defects and bifurcation defects occur. These observations indicate that the lack of one particular lobe, thinning, thickening of a specific lobe, and mis-projection of one lobe, as observed in whole brains of fly mutants (Figure 4A, B), is caused by a combination of a failure in axon branching and misguidance. Taken together, these phenotypic studies indicated that the proper composition of Dscam1 Ig 7 variants is indispensable for normal growth, branching and guidance of mushroom body axons.

We then examined how altered composition of Dscam1 isoforms affected self-avoidance of MB axonal branches. Inconsistent to previous work,(Hattori et al., 2007; Wu et al., 2012) our data demonstrate that even expression of a repertoire of up to thousands of Dscam1 isoforms (encoded within the genome of a single cell) appears not to be sufficient for normal sister branch segregation. We found that bifurcated sister branches often lose to separate normally in *Dscam1*^Δ9uD2-Δ9uD5^ mutants, as in *Dscam1*^null^ (Figure 6A, C). For example, up to 25% of *Dscam1*^Δ9uD4^ axons with bifurcated sister branches were not reliably segregated into the dorsal and medial lobes (Figure 6C). The frequency of segregation defects in *Dscam1*^Δ9uD^ mutants was not significantly less than *Dscam1*^null^ mutants (Figure 6C). However, we have observed little fascicles and crossing between sister branches in *Dscam1*^Δ9uD^ neurons (Figure 6B), while axonal sister branches in *Dscam1*^null^ neurons were extensively crossed.(Hattori et al., 2009; Hattori et al., 2007; Wang et al., 2004; Wang et al., 2002) This data indicates that the altered composition of Dscam1 isoforms did not influence the repulsion of self-branches from the same neuron. Thus, the phenotype defects observed in *Dscam1*^Δ9uD^ mutants should not be attributed to the absence of self-repulsion between sister branches from the same neuron.

**Figure 6.**
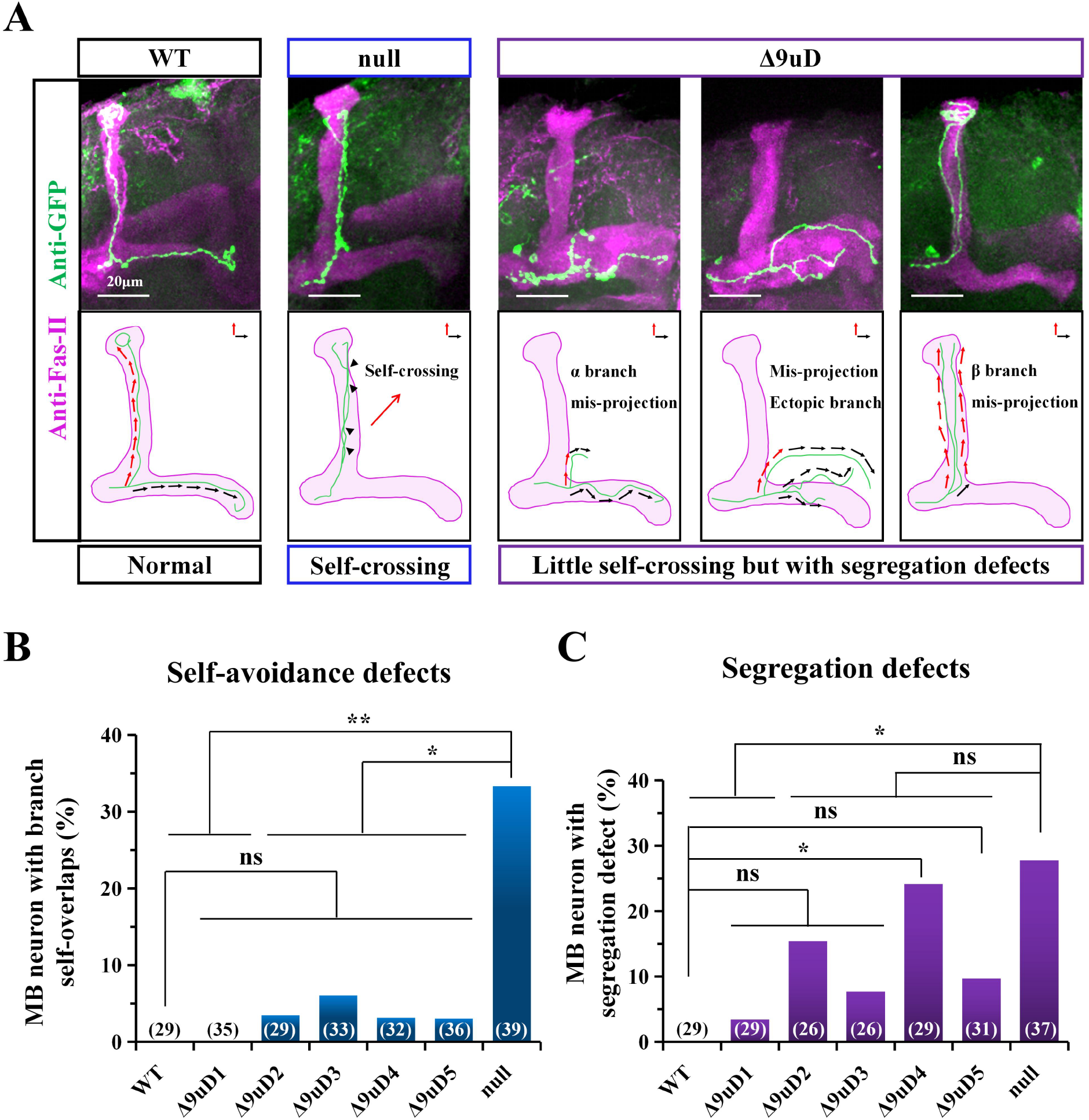
*Dscam1*^Δ9uD^ mutants exhibit abnormal branch segregation but has no effects on the self-avoidance of sister branches. **(A)** Schematic of the effects of altered isoform bias on the self-avoidance of sister branches; representative mushroom body neurons are shown. In the left panel, there are only two branches of wild type neurons extending to the apex of both lobes without self-crossing. By contrast, in the middle panel, the branches of *Dscam1*^null^ neurons frequently cross or form fascicles due to the loss of Dscam1-mediated repulsion. In the right panel, *Dscam1*^Δ9uD^ animals exhibit various types of separation defects, including one branch guides correctly, the other branch misguidance, both branches were misdirected and an ectopic branch grows from the normal branch. **(B)** *Dscam1*^Δ9uD^ mutants did not exhibit the self-avoidance defects of sister branches at a single-cell level. Numbers in parentheses represent the numbers of single-cell clone analyzed. Each image appears the crossing in 3D recorded as 1, otherwise recorded as 0. **(C)** *Dscam1*^Δ9uD^ mutants exhibited abnormal segregation of sister branches of mushroom body axons. Numbers in parentheses represent the numbers of single-cell clone analyzed which has two branches. Images which two branches project to perpendicularly lobes recorded as 0, the others recorded as 1. ns, not significant; *, P < 0.05; **, P < 0.01 (Fisher’s exact test, two-tailed).

### Altered composition of Ig 7 variants causes mild axonal defects in mechanosensory neuron

To examine a potential role of Dscam1 isoforms in the axonal patterning of MS neurons, we first compare axonal branching between adult *Dscam1*^Δ9uD1-Δ9uD5^ mutants and WT. Many aspects of *Drosophila* MS-neuron targeting are genetically hardwired, such as the position of primary or secondary axon branches, the direction of branch extension, length of branch extension, and midline crossing (Figure 7A). We found that *Dscam1*^Δ9uD1-Δ9uD5^ mutant animals had subtle to mild defects in their characteristic overall shape and the axonal branching patterns of posterior scutellar neurons, as compared with WT controls (Figure 7B, C; Figure S6A). An ipsilateral primary axon shaft was invariantly formed, but some collateral branches were lost or impaired, varying in different *Dscam1*^Δ9uD1-Δ9uD5^ mutants (Figure 7B, C; Figure S6A). For example, ∼15% of the *Dscam1*^Δ9uD2^ and *Dscam1*^Δ9uD5^ neurons completely or partially lost their lateroanterior branches (branch 2), while 100% of wild-type control clones had these branches. Further statistics analyses revealed a significant decrease in the length of branch 2 in *Dscam1*^Δ9uD2^ compared with WT control (Figure 7D). These data indicate that these differences at same position persist in the same mutants and in different genetic backgrounds (Figure 7D; Figure S6A). Because similar branching defects have been observed in mutants with the deletion of docking site of exon 6 cluster where Ig3 variant composition was altered,(Dong et al., 2022) we conclude that the defects seen in *Dscam1*^Δ9uD^ mutants derives solely from the altered composition of Dscam1 Ig 7 variants.

**Figure 7.**
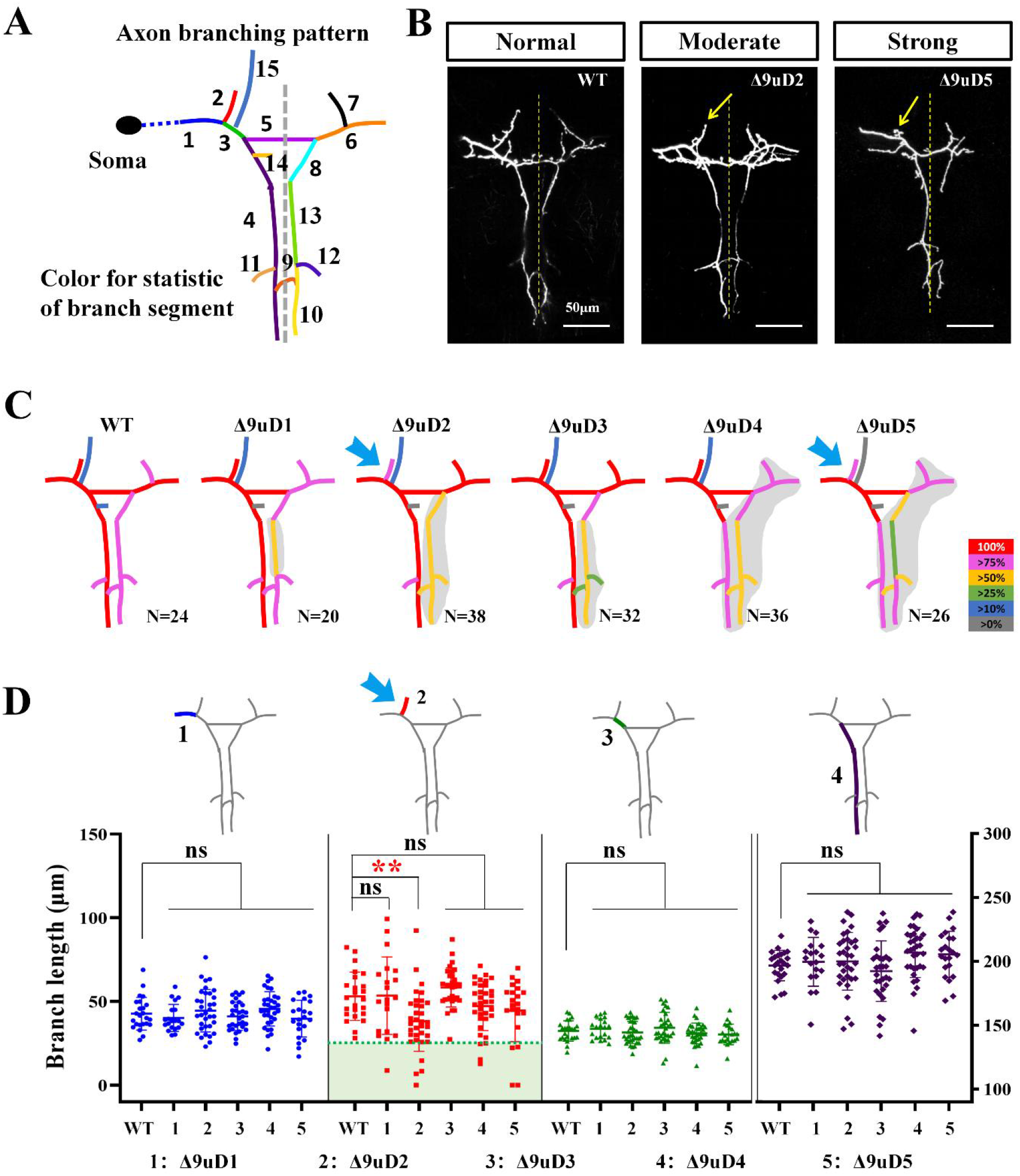
*Dscam1*^Δ9uD^ mutants exhibits subtle axonal defects in mechanosensory neuron. (see also Figure S6) **(A)** Schematic of the axon trajectory of a single mechanosensory (MS) neuron within the CNS. Different color represents a segment of statistical length and frequency. **(B)** Branching patterns of MS neurons in control and mutant animals. Yellow arrows indicate missing or shorter lateroanterior branches. The dashed line indicates the midline of the CNS; Scale bar, 50μm. **(C)** Frequency of each segment of MS neuron in wild type and *Dscam1*^Δ9uD1-Δ9uD5^ are shown. The right subscript letter N represents the number of analyzed neurons. Different color represents different frequency of segments. **(D)** Quantification of the length of segment 1 to 4 of MS neurons in *Dscam1*^Δ9uD1-Δ9uD5^. The corresponding statistical segment is shown on each cluster scatter plot. The missing branch length is denoted as 0. *, P < 0.05; **, P < 0.01; ns, not significant (Student’s t-test, two-tailed). Quantification of the length of segment 5 to 8 of MS neurons in *Dscam1*^Δ9uD1-Δ9uD5^ were showed in supporting information.

## Discussion

Our present studies reveal extensive noncanonical functions of Dscam1 isoforms in neuronal wiring. We used *cis* mutagenesis to generate mutant flies bearing normal number of Dscam1 isoforms albeit with altered isoform composition. These Dscam1 isoform composition-altered mutants exhibited normal dendritic self-avoidance and self/non-self discrimination in da neurons, which is consistent with canonical self-avoidance model. Surprisingly, these mutants exhibited strikingly distinct spectra of phenotypic defects in three classes of neurons: up to ∼60% defects in mushroom bodies, a significantly increased branching and growth in da neurons, and mild axonal branching defects in mechanosensory neuron. These data suggest that Dscam1 isoform composition is cell-autonomously required for normal growth in diverse neurons. Below, we discuss the molecular mechanisms that Dscam1 isoforms mediate wiring of MB axons and da neuron dendrites. We also discuss the importance of the specificity of individual (subsets of) Dscam1 isoforms.

### Proper Dscam1 isoform composition is required for MB axonal wiring

The canonical model of Dscam1-mediated self-avoidance has been illustrated in the axonal patterning of mushroom body neurons.(Hattori et al., 2009; Hattori et al., 2007; Hattori et al., 2008; Sanes and Zipursky, 2020; Zipursky and Sanes, 2010). However, this model alone cannot fully explain several findings in our present study. First, *Dscam1*^Δ9uD4^ allele, which potentially generates thousands of isoforms similar to WT, exhibited normal dendritic self-avoidance and self/non-self discrimination, but displayed more than 60% defects of mushroom bodies. Second, our mosaic analysis of singe *Dscam1*^Δ9uD4^ mutant neurons still displayed ∼70% of axon defects, ∼80% of which belong to growth and branching defects. Notably, ∼25% of *Dscam1*^Δ9uD5^ neurons have only a single branch (Figure 5E). These data demonstrate that the proper composition of Dscam1 isoforms are required for normal bifurcation of MB axons. Moreover, the shortened axon phenotype in *Dscam1*^Δ9uD^ mutants obviously could not be explained by Dscam1-mediated self-avoidance. In addition, *Dscam1*^Δ9uD5^ neurons exhibited abnormal branch segregation at high frequency. Taken together, our data support the notion that the canonical self-avoidance model is not the only mechanism governing MB axonal wiring.

### Dscam1 isoform composition is required for normal dendrite branching and growth of da neuron

Numerous studies demonstrate that Dscam1 functions to mediate dendrite patterning via homophilic repulsion in mechanosensory dendritic arborization (da) neurons.(Hattori et al., 2009; Hong et al., 2021; Hughes et al., 2007; Matthews et al., 2007; Soba et al., 2007) However, our present study shows that *Dscam1*^Δ9uD^ exhibit significant increases in dendritic branch number and total dendritic length without disrupting dendritic spacing or inter-neuronal repulsion. No obvious dendritic self-crossings were observed, and the overlapping between class I (vpda) and class III (v′pda) dendrites was not affected. Thus, in addition to its canonical role in dendrite self-avoidance, this study uncovers a novel function of Dscam1 in *Drosophila* PNS development, that is, Dscam1 autonomously regulates dendrite branching and growth. Similarly, using RNAi knockdown previous work demonstrated that Dscam1 is required for normal dendrite growth and branching but not for dendritic spacing and intraneuronal repulsion in fly motoneurons,(Hutchinson et al., 2014) indicating that self-repulsion in motoneurons might be mediated by other molecules or mechanisms. Taken together, these data suggest that Dscam1 isoforms play a conserved role in dendritic arbor growth and branching of both central and peripheral neurons.

Dscam1 isoform mediate dendritic branch formation possibly via a mechanism distinct from self-avoidance. In canonical model of self-avoidance, there is no requirement for specific isoforms; it is only of importance that the Dscam1 repertoire of each neuron is different from that of its neighbors.(Hattori et al., 2009) However, our data indicate that specific Dscam1 repertoire of each cell may affect the length and number of dendrite branches. Thus, specific Dscam1 repertoire of each neuron is cell-intrinsically required for normal dendrite growth and branching. Since PAK1 is not required for dendritic self-avoidance of da neuron,(Hughes et al., 2007; Matthews et al., 2007; Soba et al., 2007) and *Dscam1*^Δ9uD2^ mutants partially resemble Pak overexpression mutants exhibiting excessive branching of ddaE neurons, we speculate that Pak may be a candidate for Dscam1-mediated dendritic branching and growth.

### Are the specific (subsets of) Dscam1 isoforms important?

Previous genetic studies indicate that the specificity of individual Dscam1 isoforms may not be functionally important.(Cvetkovska et al., 2013; He et al., 2014; Kim et al., 2013; Schmucker et al., 2000; Wang et al., 2004; Wang et al., 2002; Zhu et al., 2006) However, these functional experiments using overexpression of one single isoform, knock-ins, or genomic deletions have either caused too strong or too weak defective phenotypes to unmask functional difference among individual isoforms. Alternatively, loss-of-function phenotype differences among them could easily be overridden by gain of function. In present study, we employed CRISPR/Cas9 technique to delete the intronic regulatory sequences, which allows us to assess the importance of the specificity of individual Ig7 variants. We reasoned that if individual Ig7 variant is equally functional, altered composition of exon 9 variants should not cause any phenotypic defects. Indeed, these mutants exhibited normal dendritic self-avoidance and self/non-self discrimination in da neurons. Collectively, we provide evidence supporting different roles for, if not individual Ig7 variant, subsets of Dscam1 Ig7s in regulating neuronal growth and branching.

## MATERIALS AND METHODS

### Materials: Fly Strains

All fruit flies were cultured on the standard cornmeal medium at 25°C constant temperature incubator. The fly strain *{nos-Cas9} attP2* was used for embryo microinjection to generate germline mutations.(Ren et al., 2013) *{nos-Cas9} attP2* was only used as a control (WT) for RNA-seq data. And *W*^*1118*^ was used as a control (WT) for neuron phenotypes and growth phenotypes. *if/Cyo, if/Cyo*.*GFP* and *sp/Cyo* were used as the balancer stocks to screen for mutants. The *221-GFP Gal4* line was used to drive *UAS-mCD8-GFP* expression in class I da neurons.(Grueber et al., 2003) *FRT42D* and *hsFLP,UAS-mCD8-GFP;FRT42D,tubP-Gal80/CyO;Gal4-OK107* were MARCM stocks to label single-cell of mushroom body (MB) neurons. *Dscam1*^21^ and *Dscam1*^17^ are *Dscam1*^null^ mutants and *Dscam1*^21^/*Dscam1*^17^ was used to analyze da neuron phenotypes.

### Generation of fly Dscam1 mutant alleles

We used the CRISPR-Cas9 system to delete or mutate the Docking site and selector sequence in the *Drosophila Dscam1* gene. Briefly, we constructed two sgRNA plasmids which target deletion sites and/or a donor plasmid with mutation sequences. Then, those sgRNA and donor plasmids were co-injected into *{nos-Cas9} attP2* embryos performed by UniHuaii CO., Ltd., China. Each offspring (G0) developed from the injectd embryo is crossed with *if/Cyo*.*GFP* flies. Then, we extract the genome of its female offspring (G1) and determine the mutation tubes by PCR analysis (*SteadyPure* Universal Genomic DNA Extraction Kit and 2x *Accurate Taq* Master Mix), and male offspring (G1) from mutation tubes are crossed with *if/Cyo*.*GFP* again. Next, we identify the male parent flies (G1) by PCR analysis to acquire the heterozygous mutation lines (G2) and cross each other to finally get the homozygous offspring. Homozygous offspring can be directly used to detect changes in the expression pattern of Dscam1 exon 9, whereas the growth phenotypes and the neuronal phenotypes need to be identified after five generations of backcrossing with *W*^*1118*^.

### RT-PCR

Total RNA from fly heads and different developmental periods was extracted from 30 flies using TRIzol reagent (Invitrogen). Total RNA was reverse transcription from *Dscam1* exon 10 using SuperScript III system (Invitrogen). The reverse transcription products were amplified by the PrimeSTAR DNA Polymerase (TaKaRa) using Exon9 specific primers. The RT-PCR products with an amplification cycle number of 25 and an anneal temperature of 60°C were used for RNA-seq, and the RT-PCR products with an amplification cycle number of 35 and an anneal temperature of 60°C were used for electrophoresis with 1.5% agarose gel.

### Utilization assay of exon variants

The RT-PCR products for RNA-seq were excised and gel purified using the Biospin Gel Extraction Kit (BioFlux). Purified products were pooled on Illumina MiSeq platform (Illumina, San Diego) according to the standard protocol by G-BIO.We calculated the relative expression levels of *Dscam1*exon 9s according to the method as previously described.(Hong et al., 2021)

### Western blot analyses

Total protein was extracted from 30 fly heads using strong RIPA lysis buffer (CW-BIO) and the protease inhibitor PMSF (Beyotime). Equal amounts of WT and mutant’s protein samples were separated by 10% Precast-Glgel Tris-Glycine PAGE gel (Sangon Biotech) and then transferred onto PVDF membranes. The membranes were blocked with 5% NON-Fat Powdered Milk (Sangon Biotech) at room temperature for 1 hour and transferred to the diluted primary antibodies to Dscam1 (ab43847, diluted 1:5000) or β-actin (ab8227, diluted 1:10,000) and incubated at 4°C overnight. After three washes with TBST, the membranes were incubated with the secondary antibodies (Goat Anti-Rabbit IgG, 1:10,000, CW-BIO) at room temperature for two hours. The immunoreactive bands of Dscam1 and β-actin protein were detected using the eECL Western Blot Kit (CW-BIO) and Tanon 5200. The Image J was used to perform grayscale analysis on the bands of Dscam1 and β-actin.

### Growth and development detection

We collect 150 virgin female and male flies each, and mix them for two days in a bottle containing a solidified juice tray and yeast extract. Replace the juice tray with a new one every three hours for a total of five times. Collect 200 embryos from each replacement juice tray and place them in a new large juice tray. After 48 hours, count the number of hatched embryos to obtain the hatching rate. 24 hours later, 30 second-instar larvae are collected from the large juice tray into the food tube, and two tubes were collected in each large juice tray. After 3-4 days, we count the number of pupae on the tube wall to obtain the pupation rate. Finally, the eclosion rate after complete eclosion was counted.

### Crawling distance detection

We collect 30 middle third instar larvae, larvae crawled freely on fruit juice medium (1:2, 3% agarose gel) and we photograph their trackway in 1 minute. Then, the trackway length was calculated by Image J.

### Climbing ability detection

We collect 90 one-day-old female and male flies each and put them into the tube with a mark of 2 cm height, 30 female or male flies per tube. Hit the tube hard so that all fruit flies fall to the bottom, and count the number of flies that climb over two-centimeter height after 10 seconds of stopping the impact. Repeat this process ten times and pause for 1 minute between each repetition. Take the average of ten trials as the final result of one tube. Finally, the climbing ability was shown by a percentage of the total number of flies in each tube.

### Immunostaining

We use third-instar larvae on the tube wall to detect the growth, branching, and coexistence of da neurons. Specifically, the dissection is performed in a silicone-coated dish containing phosphate buffered saline (PBS), epidermis were stretched and fixed with 4% paraformaldehyde at room temperature for 25 mins. The 221-GFP Gal4 stock was used to drive GFP expression in class I da neuron, HRP antibody (Cy3-conjugated Affinipure Goat Anti-Horseradish Peroxidase, diluted 1:200) was used to stain all da neurons. The larval epidermis were blocked in 5% BSA (diluted in PBST, PBS containing 0.1% Triton X-100) after three washes in PBST (20 mins each time) and incubated in HRP antibody at 4°C overnight. After three washes in PBST for 20 mins each, samples were mounted in prolong gold antifade reagent (Invitrogen). The immunofluorescence staining was imaged by confocal microscope LSM800 (Carl Zeiss).

Dissecting the heads of *Drosophila* within 5 days after eclosion for mushroom body immunofluorescence staining. The dissection is performed in cold PBS and fixed in 4% paraformaldehyde for 45 mins at room temperature. After three washes by PBST, 20 mins each time, removing the white tissue on the surface of the brain on a glass slide and then blocked in 5% BSA for 1h at room temperature. Three washes again, the brains were incubated with the primary antibodies (anti-FasII, DSHB, diluted 1:2 in PBST) for 36-48 hours at 4°C. After the standard washing, the brains were incubated with the secondary antibodies (Alexa594-goat-anti-mouse IgG, Earthox, diluted 1:400 in PBST) for 5 hours at room temperature. Wash three times again, the brains were mounted in prolong gold antifade reagent (Invitrogen). The immunofluorescence staining was imaged with a laser scanning confocal microscope LSM800 (Carl Zeiss).

### Induction and phenotypic analysis of MARCM clones

Single-cell clones of mushroom body neurons of *Dscam1* mutants were generated using MARCM techniques.(Lee and Luo, 1999) First, female *Dscam*^*Δ9*uD^ mutants were hybridized with *FRT42D* lines to produce recombinant mutants with FRT sites. Next, Recombinant mutants males mate with *hsFLP,UAS-mCD8-GFP;FRT42D,tubP-Gal80/CyO;Gal4-OK107* to generate GFP expression clones. Then, mitotic recombination was induced via heat shock, the heat shock condition is 37 °C water bath for 45 mins, 25 °C incubation for 15 mins, and 37 °C water bath for 45 mins again. Immunostaining of adult brains was performed as described above.

### Mechanosensory neuron analysis

We fixed the flies and removed the bristles of their dorsal plate pSC neurons and then fixed them in 4% paraformaldehyde overnight. DiD and DiI (20 mg/mL) were infused into two pSC neuron pores, respectively. Next, flies were immersed in the 0.2M pH9.5 CB buffer for 1-5 days. Finally, ventral nerve cord (VNC) were dissected and imaging was performed by confocal microscope LSM800 (Carl Zeiss).

### Statistical analysis

Quantitative analysis was performed from three biological replicates. Error bars represent mean±SD. The significance of the difference was determined by the two-tailed Student’s t-test, and ns p > 0.05, * P < 0.05, ** P < 0.01, and *** P < 0.001 were used to indicate statistical significance. Neuron length is calculated by neuron J.

## Supplementary information

Supplementary Figs. 1–6

## Acknowledgments

This work was supported by research grants from the National Key Research and Development Program of China (2021YFE0114900), the National Natural Science Foundation of China (91940303, 91740104), the Natural Science Foundation of Zhejiang Province (LD21C050002), and the Starry Night Science Fund at Shanghai Institute for Advanced Study of Zhejiang University (SN-ZJU-SIAS-009).

## Author contributions

YJ conceived of this project. SZ, XY, HD, JZ and LW designed and performed the experiments; SZ, LW, BX, HD, PG, YZ and JZ participated in the construction and screening of the mutants; SZ, XY, LL, YF, YD performed phenotype analysis. SZ, PG, GL and FS analyzed the data; YJ, HH, JH, SZ, and GL wrote the manuscript; all authors discussed the results and commented on the manuscript.

