## Supplemental File for "*Cis* mutagenesis *in vivo* reveals extensive noncanonical functions of Dscam1 isoforms in neuronal wiring"

[1]MOE Laboratory of Biosystems Homeostasis & Protection and Innovation Center for Cell Signaling Network, College of Life Sciences, Zhejiang University, Hangzhou, Zhejiang, ZJ310058, [2]Department of Neurosurgery and State Key Laboratory of Biotherapy, West China Hospital, Sichuan University, Chengdu 610041, [3]Institute of Insect Sciences, Zhejiang University, Hangzhou, Zhejiang, ZJ310058, China, PR China.

### Supplementary information

Supplementary Figs. 1–6

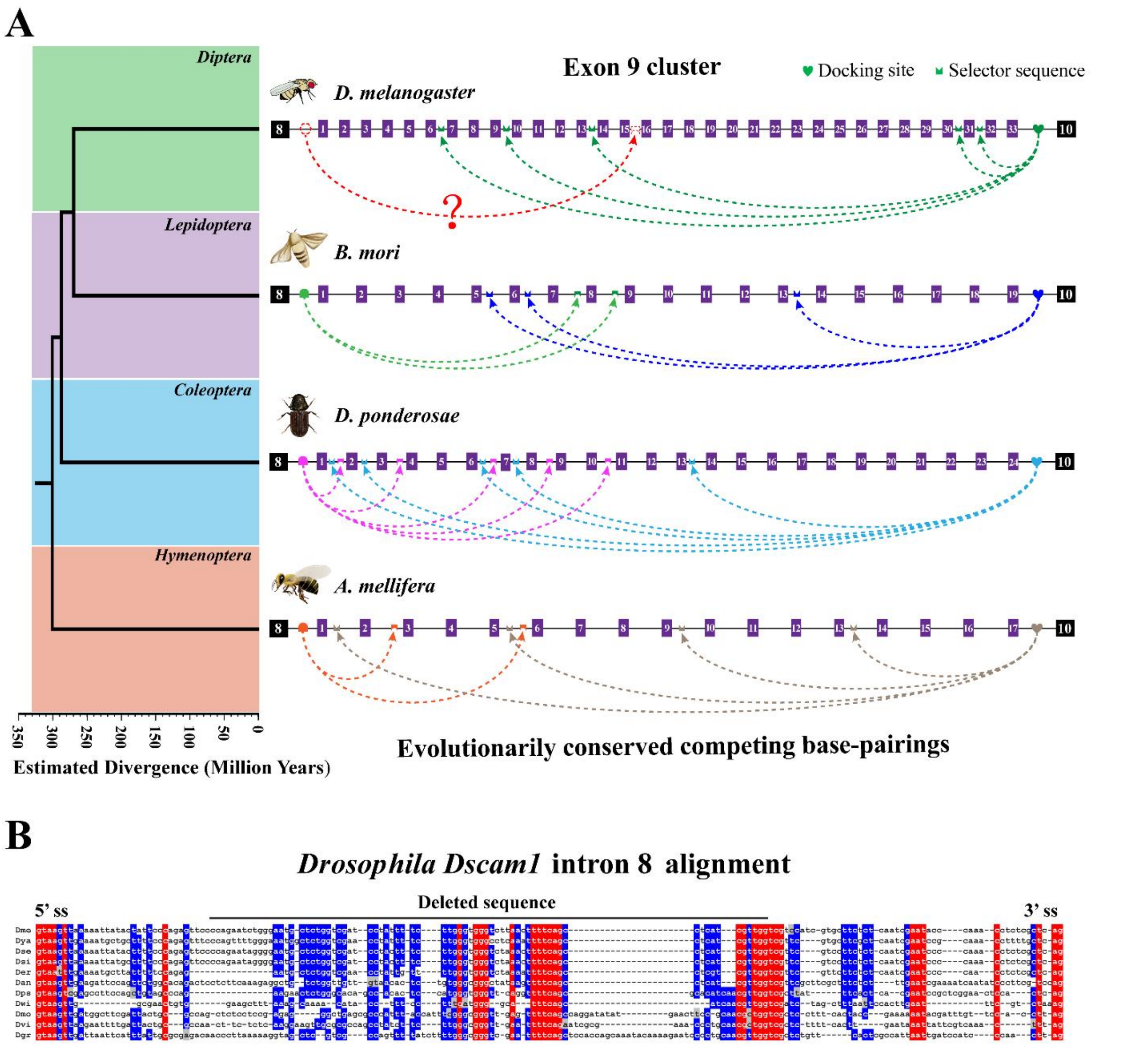


**Figure S1. Conserved base pairings in exon 9 cluster of insects *Dscam1*. (Related to Figure 1)**

**(A)** A phylogeny of conserved base pairings in governing alternative splicing of exon 9 clusters during insect evolution. Exons, docking sites and selectors are not drawn to scale. Sequence-specific docking sites and selector sequences are shown in different colors. The dashed arrow represents base pairing interaction between the docking site and selector sequence. Inter-intronic base pairings in *D. melanogaster* was confirmed.(Hong et al., 2021) The base pairings between the docking site and selector sequence for exon 9 cluster of *Hymenopteran Dscam1* were experimentally verified.(Yue et al., 2016) The base pairings between the docking site and selector sequence in lepidopteran *P. xylostella*, and Coleopteran *T. castaneum* were predicted by comparative genome comparison and structural modeling.(Dong et al., 2021) These data showed that competing base pairings are conserved in exon 9 cluster of insect *Dscam1*, but the docking site and selector sequence are clade-specific.

**(B)** Alignment of intron 8 sequence of *Drosophila Dscam1*. The invariant nucleotides at each position are shaded in red, while the most identical nucleotides are shaded in blue.

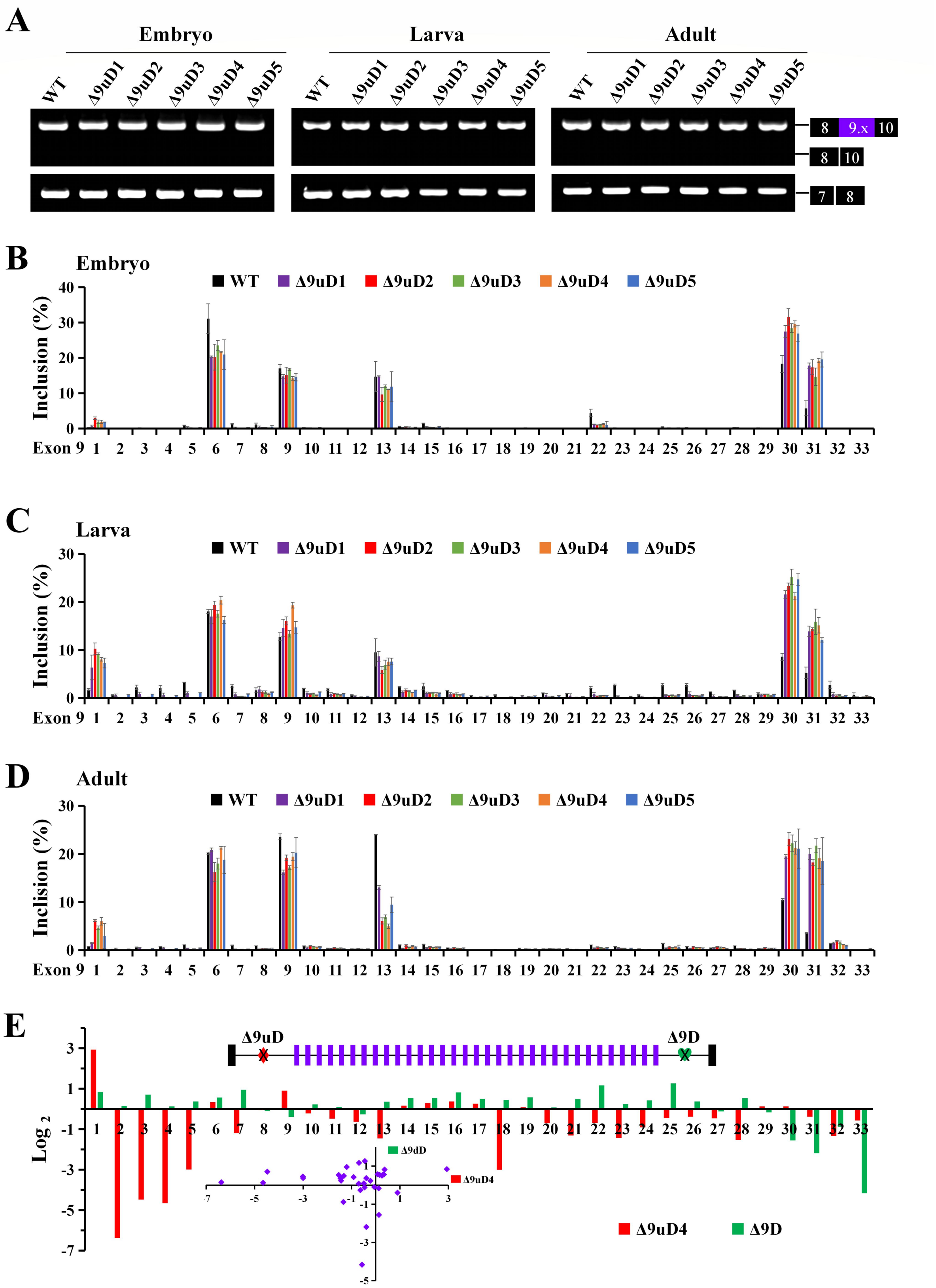


**Figure S2. Expression pattern of the exon 9 inclusion in various developmental stages and tissues of mutants. (Related to Figure 1)**

**(A)** RT-PCR analysis showed that the inclusion of exon 9 is not affected in various developmental stages.

**(B-D)** *Dscam1*$^{\Delta 9uD}$ show similar change expression at different developmental stages.

**(E)** The $\log_2$ fold change of the frequency of variable exon 9 inclusion in *Dscam1*$^{\Delta 9uD4}$ was largely contrast with the change pattern in *Dscam1*$^{\Delta 9D}$, which lack the downstream docking site.(Hong et al., 2021)

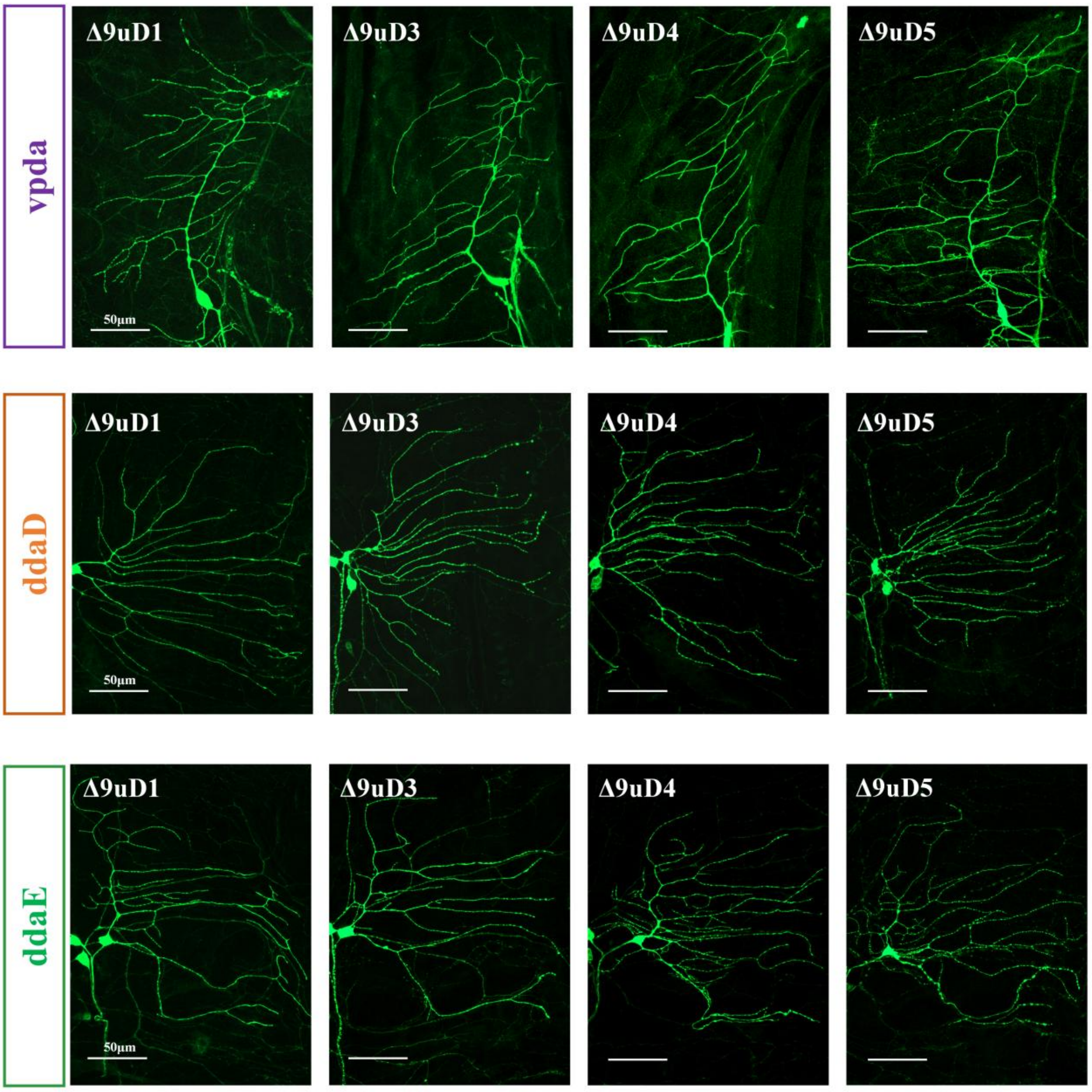


**Figure S3. Class I da neurons exhibit dendritic patterning defects. (Related to Figure 2, 3)** Representative images of the Class I da neuron including vpda, ddaD and ddaE in different *Dscam1* mutants. All neurons were visualized with GFP. Scale bar, 50μm.

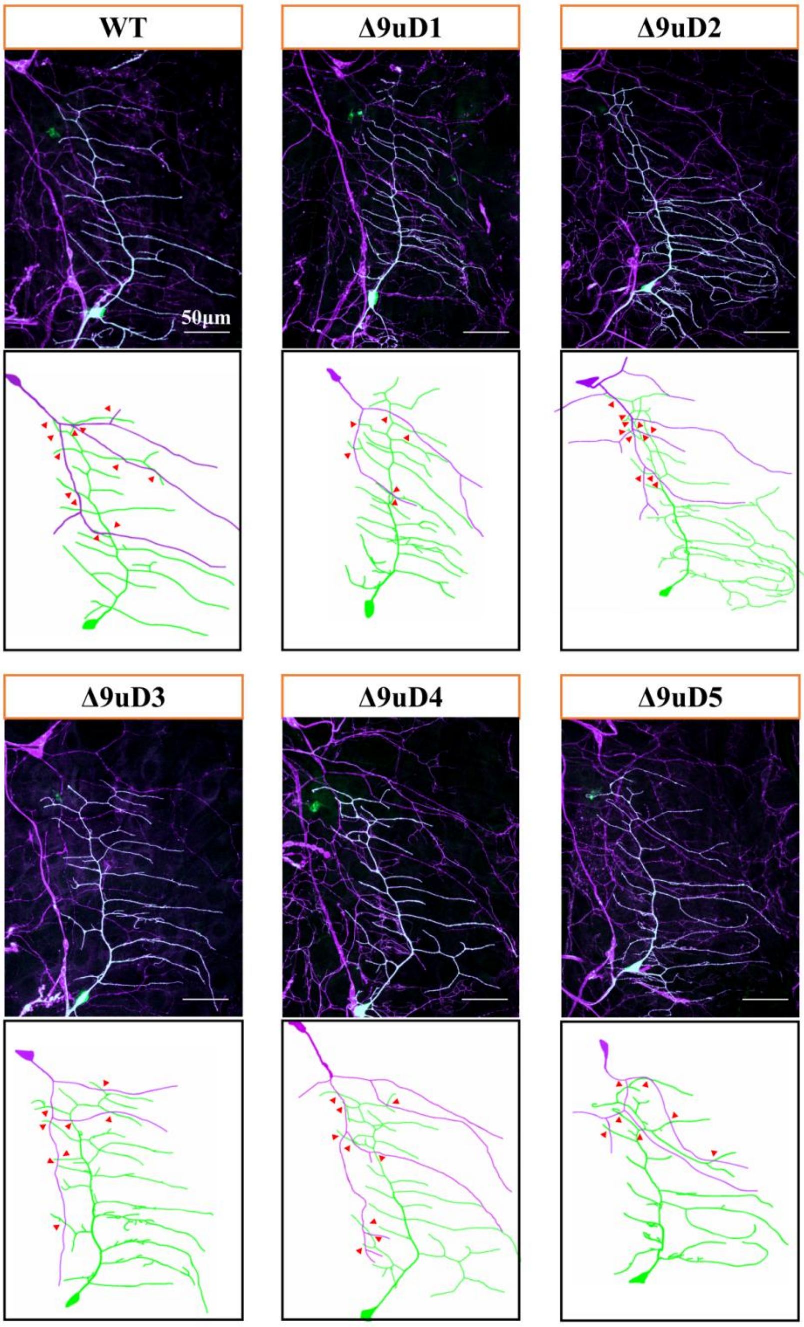


**Figure S4. The number of overlaps between class I and class III dendrites were indistinguishable between *Dscam1*$^{\Delta 9uD1\text{-}5}$ mutants and wild-type controls. (Related to Figure 2)** The directions of the class I (green) and class III (magenta) branches are depicted under each representative image. Red arrowheads directing the overlaps of class I and class III. Scale bar, 50μm.

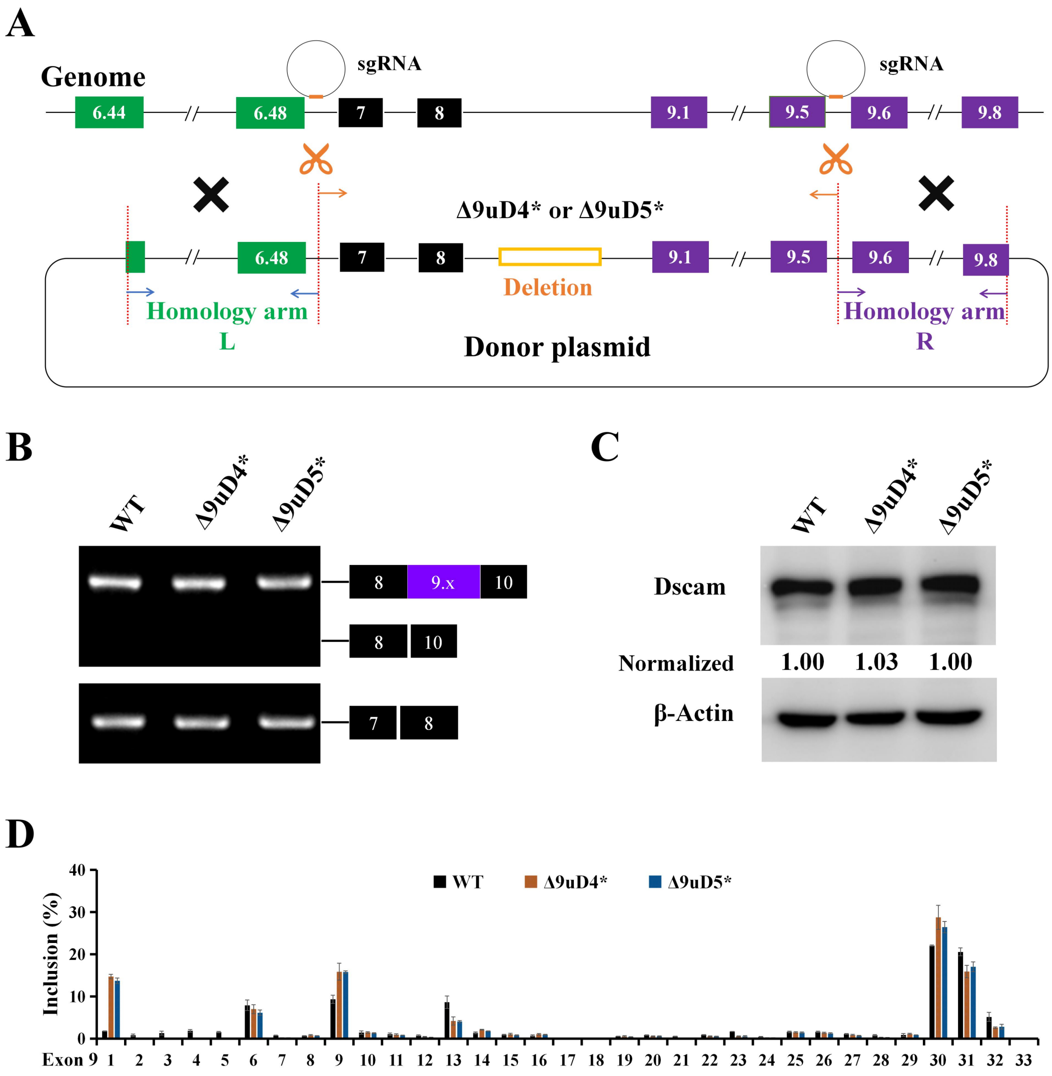


**Figure S5. Construction and molecular characterization of *Dscam1*$^{\Delta 9uD*}$ mutants. (Related to Figure 4)**

**(A)** Construction of *Dscam1*$^{\Delta 9uD4*}$ and *Dscam1*$^{\Delta 9uD5*}$ homologous recombination mutants by CRISPR/Cas9 system. The location of sgRNAs are shown. The sequence of *Dscam1*$^{\Delta 9uD4*\text{-}\Delta 9uD5*}$ and *Dscam1*$^{\Delta 9uD4\text{-}\Delta 9uD5}$ are identical.

**(B)** RT-PCR analysis showed that the overall inclusion of exon 9 is not affected in *Dscam1*$^{\Delta 9uD4*}$ and *Dscam1*$^{\Delta 9uD5*}$.

**(C)** Western blotting analysis show that the Dscam1 level of *Dscam1*$^{\Delta 9uD*}$ are same as wild type.

**(D)** *Dscam1*$^{\Delta 9uD4*}$ and *Dscam1*$^{\Delta 9uD5*}$ show similar Exon 9 splicing pattern with *Dscam1*$^{\Delta 9uD4}$ and *Dscam1*$^{\Delta 9uD5}$.

**A** **Frequency of occurrence of each branch of MS neuron in different genotypes**

| Genotypes | N | Percentage of each Branch (%) | | | | | | | | | | | | | | |
|---|---|---|---|---|---|---|---|---|---|---|---|---|---|---|---|---|
| | | 1 | 2 | 3 | 4 | 5 | 6 | 7 | 8 | 9 | 10 | 11 | 12 | 13 | 14 | 15 |
| *WT* | 24 | 100 | 100 | 100 | 100 | 100 | 87.5 | 100.0 | 95.8 | 79.2 | 91.7 | 95.8 | 83.3 | 79.2 | 16.7 | 16.7 |
| *Δ9uD1* | 20 | 100 | 100 | 100 | 100 | 100 | 95.0 | 95.0 | 80.0 | 95.0 | 90.0 | 95.0 | 80.0 | 60.0 | 10.0 | 15.0 |
| *Δ9uD2* | 38 | 100 | 89.5 | 100 | 100 | 100 | 100 | 100 | 68.4 | 63.2 | 94.7 | 73.7 | 57.9 | 57.9 | 5.3 | 21.1 |
| *Δ9uD3* | 32 | 100 | 100 | 100 | 100 | 100 | 100 | 100 | 87.5 | 50.0 | 78.1 | 62.5 | 40.6 | 59.4 | 6.3 | 12.5 |
| *Δ9uD4* | 36 | 100 | 100 | 100 | 97.2 | 100 | 97.2 | 97.2 | 83.3 | 75.0 | 88.9 | 75.0 | 61.1 | 66.7 | 2.8 | 11.1 |
| *Δ9uD5* | 26 | 100 | 88.5 | 100 | 92.3 | 100 | 88.5 | 92.3 | 69.2 | 69.2 | 88.5 | 76.9 | 69.2 | 42.3 | 3.8 | 3.8 |

**B**

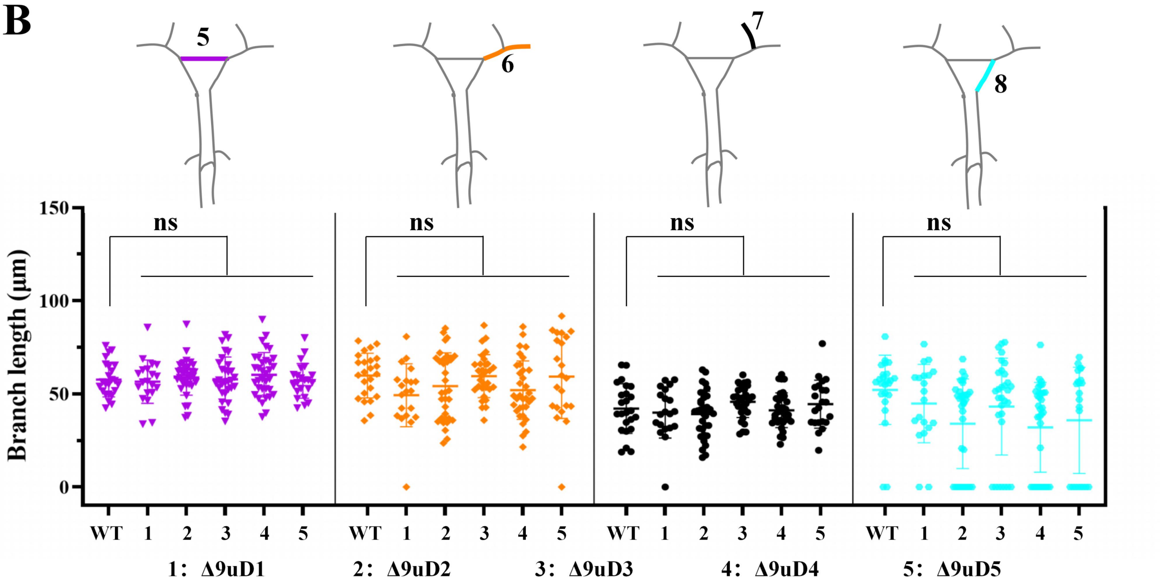


**Figure S6. Branching patterns of posterior scutellar neurons in different mutants. (Related to Figure 7)**

**(A)** A table of average branching patterns in each mutant and control.

**(B)** Quantification of the length of segment 5 to 8 of MS neurons in *Dscam1*$^{\Delta 9uD1\text{-}5}$. The corresponding statistical segment is shown on each cluster scatter plot. The missing branch length is denoted as 0. ns, not significant (Student's t-test, two-tailed).

### 参考文献

Dong, H., Li, L., Zhu, X., Shi, J., Fu, Y., Zhang, S., Shi, Y., Xu, B., Zhang, J., Shi, F., *et al.* (2021). Complex RNA Secondary Structures Mediate Mutually Exclusive Splicing of Coleoptera Dscam1. Front Genet *12*, 644238.

Hong, W., Zhang, J., Dong, H., Shi, Y., Ma, H., Zhou, F., Xu, B., Fu, Y., Zhang, S., Hou, S., *et al.* (2021). Intron-targeted mutagenesis reveals roles for Dscam1 RNA pairing architecture-driven splicing bias in neuronal wiring. Cell Reports *36*.

Yue, Y., Yang, Y., Dai, L., Cao, G., Chen, R., Hong, W., Liu, B., Shi, Y., Meng, Y., Shi, F., *et al.* (2016). Long-range RNA pairings contribute to mutually exclusive splicing. RNA *22*, 96-110.
